# CYP46A1 Activation Improves Retinal Neovascularization in a Mouse Model of Retinopathy of Prematurity

**DOI:** 10.1101/2025.10.14.682405

**Authors:** Briah Bailey, Josephine Rudd Zhong Manis, Gayatri Seth, Shubhra Rajpurohit, Allston Oxenrider, Pamela M. Martin, Ravirajsinh N. Jadeja, Menaka C. Thounaojam

## Abstract

Previous studies have shown the metabolic and regulatory significance of CYP46A1 in the adult retina; however, its role in the developing retina is unknown. Here, we evaluate CYP46A1 expression and the impact of its activation in the developing mouse retina under normal and pathological conditions. Seven-day-old (P7) C57BL/6J mice maintained in room air (controls) or subjected to oxygen-induced retinopathy (OIR) were treated with/without 20 mg/kg efavirenz (EFV), a CYP46A1 activator administered intraperitoneally from P7 to P17. Retinal cross sections and flat mounts were prepared to study retinal vasculature morphology, Müller and microglia activation, and ganglion cell viability. EFV treatment significantly reduced pathological neovascularization and the size of avascular and hypoxic areas in OIR mice retinas. EFV treatment additionally limited reactive gliosis and microglia activation and improved retinal ganglion cell survival in OIR mice. The current study demonstrates the developmental regulation of CYP46A1 and the dysregulated expression and levels of the downstream metabolite 24-Hydroxycholesterol (24HC) in OIR mice. The study further suggests that pharmacological CYP46A1 activation may improve key pathological features associated with pathological neovascularization in OIR mice.

## 1. Introduction

Retinopathy of prematurity (ROP), an ocular complication of prematurity, is a leading cause of childhood blindness [1-3]. Low birth weight (less than 1250 grams), low gestational age (less than 31 weeks), and prolonged oxygen therapy are consistently associated with increased disease risk [1-3]. To date, therapies for ROP are applied in the most advanced, proliferative stages of the disease and essentially involve ablation of the peripheral retina through surgical procedures like laser photocoagulation, cryotherapy, and vitrectomy [4-6]. Intravitreal injections of anti-vascular endothelial growth factor (VEGF) are also commonly applied to halt retinal neovascularization (RNV) [4-6]. However, this treatment may also block the normal vascular development still taking place in the premature infant retina. Therefore, even when treated, children affected by ROP may still experience substantial vision loss and/or are more likely than others to develop visual impairments later in life (e.g., nearsightedness, strabismus, amblyopia, and glaucoma) [7-10]. Therefore, identifying novel mechanisms and developing new therapies for ROP is important.

Bile acids have significant potential with respect to limiting RNV [11-13]. However, the underlying mechanisms are not fully understood. Bile acids are essential for cholesterol catabolism and cytochrome P450 (CYP) enzymes are critically involved. Several studies have implicated roles for CYP enzymes in the regulation of retinal angiogenesis [14, 15], however whether they are expressed and function similarly in the premature retina is not known. The present study investigates the expression of CYP27A1 and CYP46A1 in the developing mammalian retina in the context of pathologic angiogenesis using oxygen-induced retinopathy (OIR) mice. CYP27A1 functions as a sterol 27-hydroxylase, whereas CYP46A1 is responsible for cholesterol 24-hydroxylation [16, 17]. Both CYPs play crucial roles in removing cholesterol by generating 24-hydroxycholesterol (24HC) and 27-hydroxycholesterol (27HC), oxysterols that facilitate the movement of tissue cholesterol to the systemic circulation [18-20]. 24HC and 27HC also interact with liver X receptors (LXRs) [21], and serve as substrates for the synthesis of primary bile acids in the retina [22], which is important given the protective signaling elicited by bile acids in retinal diseases, including ROP [13].

The current study is the first to demonstrate CYP27A1 and CYP46A1 in the normal developing mammalian retina. Further, our findings expose a novel regulatory role for CYP46A1 in the pathologic angiogenesis characteristic of OIR/ROP and establish CYP46A1 as a therapeutic target for mitigating retinal inflammation and neovascularization for improved ROP management.

## 2. Materials and Methods

### 2.1. Oxygen-Induced Retinopathy (OIR) Model

All animal procedures adhered to Association for Research in Vision and Ophthalmology guidelines and were approved by the Meharry Medical College Institutional Animal Care and Use Committee. C57BL/6J mice (Jackson Laboratories, Bar Harbor, ME, USA) were housed under standard conditions with a 12-hour light/dark cycle and ad libitum access to food and water.

OIR was induced following the Smith et al. protocol, previously validated in our studies [11, 23, 24]. On postnatal day 7 (P7), pups and their dam were placed in a hyperbaric chamber (BioSpherix, Parish, NY, USA) with iris ports enabling i*n vivo* manipulation without disrupting oxygen levels. Oxygen was maintained at 70% from P7 to P12 to induce central vaso-obliteration. Mice were then returned to room air (21% oxygen) until P17 to promote retinal neovascularization (RNV). Age-matched controls were kept in room air from P0 to P17.

### 2.2. Efavirenz (EFV) Administration

EFV (Cayman Chemical, Ann Arbor, MI, USA) was dissolved in 0.01% DMSO and administered intraperitoneally at 20 mg/kg/day from P7 to P17. Vehicle controls received 0.01% DMSO alone. Eyes were harvested at P17 for downstream analyses.

### 2.3. Retinal Vascular Quantification

At P17, eyes were fixed in 4% paraformaldehyde and retinas dissected for flat mount preparation. Vessels were stained using biotinylated Isolectin GS-IB4 (0.2 mg/ml; Invitrogen, Carlsbad, CA, USA) and Texas Red–conjugated avidin D overnight at 4°C. Imaging was performed using a Zeiss LSM 780 Inverted Confocal microscope. Quantification of vaso-obliteration and neovascularization was automated using the OIRSeg tool (http://oirseg.org). Angiogenesis parameters were further analyzed using AngioTool software (NIH/NCI).

### 2.4. Western Blot Analysis

Retinal lysates were prepared at P7, P12, P14, P17, and P23 using RIPA buffer (Thermo Fisher, Waltham, MA, USA) supplemented with 1% phosphatase and protease inhibitors (Sigma-Aldrich, St. Louis, MO, USA). Protein (30–50 µg) was resolved by SDS PAGE and transferred to PVDF membranes. Membranes were blocked with 5% skim milk and probed with primary antibodies: CYP46A1 (1:1000, RayBiotech), CYP27A1 (1:1000, Abcam), RBPMS (1:500, GeneTex), GFAP (1:1000, Abcam), VCAM-1 (Abclonal), and ICAM-1 (1:1000, Santa Cruz Biotechnology). β-actin (1:3000, Sigma-Aldrich) was used as a loading control. Secondary detection was performed using chemiluminescence (Thermo Fisher) and Azure 300 Imaging System. Band intensities were quantified using NIH ImageJ software and expressed as fold change relative to controls.

### 2.5. Immunofluorescence

Retinal cryosections and flat mounts were fixed in 4% paraformaldehyde, blocked with 1X Power Block (Biogenex), and incubated overnight at 4°C with primary antibodies: CYP27A1 (1:100, Abcam), CYP46A1 (1:100, Abcam), RBPMS (1:500, GeneTex), GFAP (1:100, Abcam), and IBA-1 (1:100, FUJIFILM Wako). After washing with 0.1% Triton X-100 in PBS (pH 7.4), slides were incubated with fluorescent secondary antibodies (Life Technologies) and mounted with DAPI-containing fluoroshield (Sigma-Aldrich). Images were acquired at 20X magnification using a Zeiss Axioplan-2 fluorescence microscope.

### 2.6. RNA Sequencing

Total RNA was extracted from P17 retinas using the RNeasy kit (Qiagen). cDNA libraries were generated via strand-specific amplification and sequenced using paired-end 50-bp reads on an Illumina HiSeq 2500 platform at Augusta University’s Genomics Core. Data were analyzed using Tophat2 and Cufflinks, and heatmaps were generated to visualize differential gene expression in sterol metabolism pathways.

### 2.7. ELISA for 24S-Hydroxycholesterol

Retinal homogenates from RA, OIR, and EFV-treated OIR mice were analyzed using the 24S-Hydroxycholesterol ELISA Kit (Abcam, ab204530). Samples were incubated in wells pre-coated with goat anti-rabbit IgG, followed by biotinylated 24-OHC and streptavidin-HRP. After TMB substrate addition, absorbance was measured at 450 nm. Concentrations were calculated from a standard curve and expressed as ng/g retina.

### 2.8. Statistical Analysis

Data are presented as mean ± SEM from 3-7 biological replicates. Statistical analyses were performed using the Kruskal–Wallis test for multiple group comparisons and unpaired t-tests for pairwise comparisons, with statistical significance set at p < 0.05. Graphs were generated using GraphPad Prism v10 (GraphPad Software, San Diego, CA, USA).

## 3. Results and Discussion

Retina intricately balances cholesterol synthesis and output pathways to avoid the detrimental effects of excess tissue cholesterol [25, 26]. Cholesterol conversion to oxysterols (27-hydroxycholesterol, 27HC; 5-cholestenoic acid, 27COOH; 7α-hydroxy-3-oxo-4-cholestenoic acid, 7HCA; and 24-hydroxycholesterol, 24HC) by cytochrome P450 enzymes 27A1 and 46A1 (CYP27A1 and CYP46A1, respectively) is key to elimination [25, 26]. Further, 27HC and 24HC interact with LXR [27] and serve as substrates for the synthesis of primary bile acids in retina. The importance of CYP27A1 and CY46A1 in retina is highlighted further by the demonstration of defective cholesterol clearance/increased lipid deposition and retinal neovascularization in adult mice lacking CYP27A1 and CYP46A1 [16, 27]. However, the expression and function of these enzymes in the developing retina and ROP remains unknown.

The mammalian retina continues to develop postnatally. Understanding key metabolic and molecular changes that occur pre- and postnatally in health and disease, the purpose of the present study, may provide critical information toward the development of new therapies for neonatal retinal diseases. We and others have demonstrated the therapeutic potential of bile acids in retinal neovascularization [11, 13, 22]. However, the underlying mechanisms to explain these effects are not fully understood. RNA sequencing analyses (Fig. 1A-C) performed using retinal samples obtained from RA and OIR mice at P17 confirmed the alteration of key players in sterol and BA synthesis pathways in OIR mice. CYP46A1 was notably downregulated compared to RA controls (Fig. 1B), as confirmed by Western blotting of CYP46A1 expression (Fig. 1D) and ELISA analyses for levels of 24HC (Fig. 1E). Interestingly, given the surrogate relationship between CYP27A1 and CYP46A1 [19], RNA sequencing studies did not reveal significant alterations in CYP27A1. However, because little is known regarding the expression and functional relevance of these enzymes in the normal developing retina, we evaluated their expression in P7-P23 mouse retinas. The expression of both CYPs was evident by P12 and increased progressively with age (Fig. 2A-B). However, congruent with RNA sequencing data above, only CYP46A1 expression was disrupted significantly in OIR (Figs. 2C-D). As such, the remainder of the study focuses on CYP46A1 expression and function in OIR.

**Figure 1.**
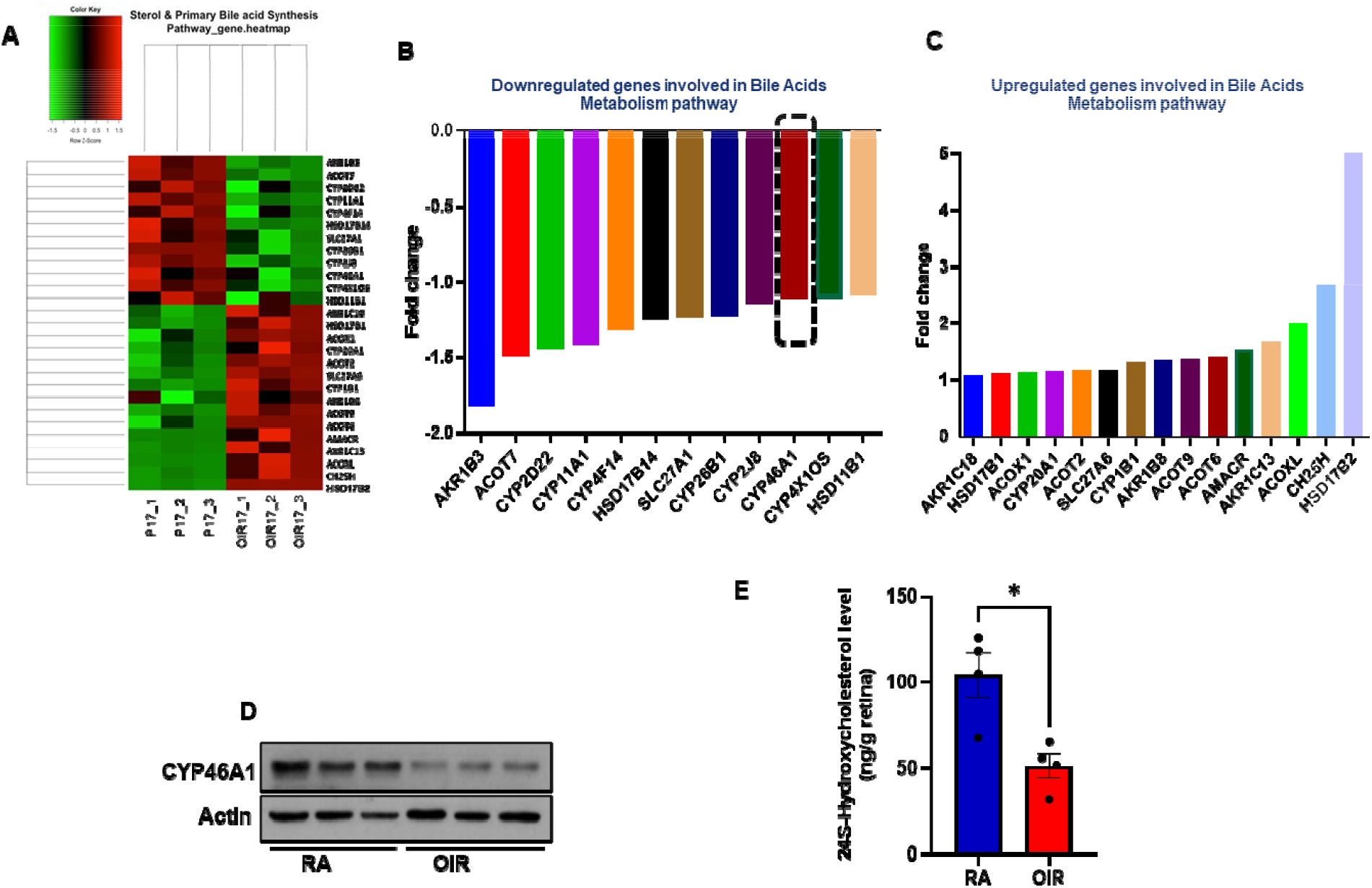
RNA sequencing was performed using total RNA extracted from room air (RA) and oxygen-induced retinopathy (OIR) mouse retinas at postnatal day 17 (P17). (A) Heat map illustrating genes up-(red) or down-regulated (green) in sterol metabolism pathways, expressed as fold-changes in gene expression in (B-C). Results are shown as mean for n = 3 retinas per group; each obtained from a different mouse. (D) Western blot evaluation of CYP46A1 protein expression in RA and OIR mice. Values are presented as mean ± SEM (*n*=3 retinas per group, each obtained from a different mouse). (E) ELISA quantification of 24S-hydroxycholesterol in retinal samples showing changes in 24S-Hydroxycholesterol levels in OIR mice. Values are presented as mean ± SEM (n = 4 retinas per group, each obtained from a different mouse). *p<0.05 vs. RA or OIR.

**Figure 2.**
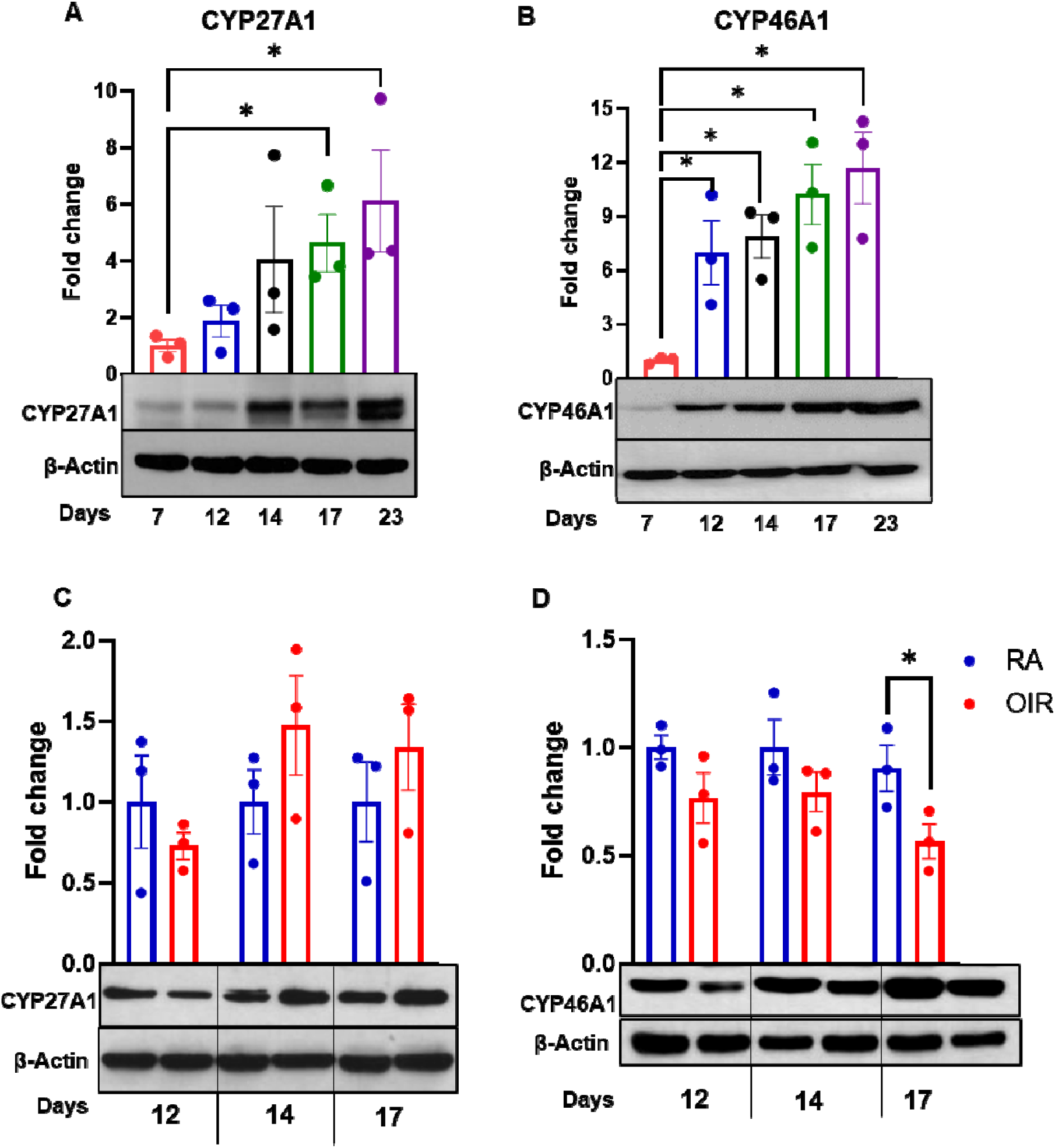
(A-B) Western blot evaluation of CYP27A1 and CYP46A1 protein in the developing mouse retina (postnatal day 7 to 23), or (C-D) in mice maintained in room air (RA) or subjected to oxygen-induced retinopathy (OIR) at P12, 14 and 17. Results were normalized to β-actin and expressed as fold change; mean ± SEM (n = 3 retinas per group, each obtained from a different mouse). *p<0.05 vs. postnatal day 7 (P7) or RA.

Because CYP46A1 expression was found to be downregulated in OIR, we next evaluated the impact of CYP46A1 expression in OIR pathology in the presence or absence of the CYP46A1 activator efavirenz (EFV). EFV, an FDA-approved antiretroviral medication for HIV treatment, has also been tested in clinical trials in patients with mild cognitive impairment or early dementia due to Alzheimer’s disease [28]. EFV activates CYP46A1 by binding to the allosteric site on the CYP46A1 surface [29]. In OIR mice treated with 20 mg/kg EFV, a dose derived from prior studies of its use therapeutically [30], from P7-P17 (both hyperoxia and hypoxia phases), the size of the avascular area and extent of pathological neovascularization was significantly reduced (Figs. 3A-D). Further, AngioTool quantification of vessel morphometric and spatial parameters revealed that compared to RA mice, total vessel length and the number of vessel junctions were significantly lower in OIR mice, whereas the number of vessel endpoints was significantly increased (Fig. 3E-G). However, in EFV-treated OIR mice, total vessel length and the number of vessel junctions improved, and the number of vessel end points decreased (Fig. 3E-G). To better understand the effect of EFV treatment on CYP46A1 activity in the retina, we analyzed 24-OHC levels in RA, OIR, and EFV-treated OIR mice retinas using an ELISA assay. As shown in Fig. 3H, levels of 24-OHC were significantly improved in EFV-treated mice retinas compared to OIR samples, providing evidence for increased CYP46A1 activity. Collectively, these data provide evidence in support of the positive influence of maintaining CYP46A1 expression against the development of OIR pathology in mice and potentially ROP in humans.

**Figure 3.**
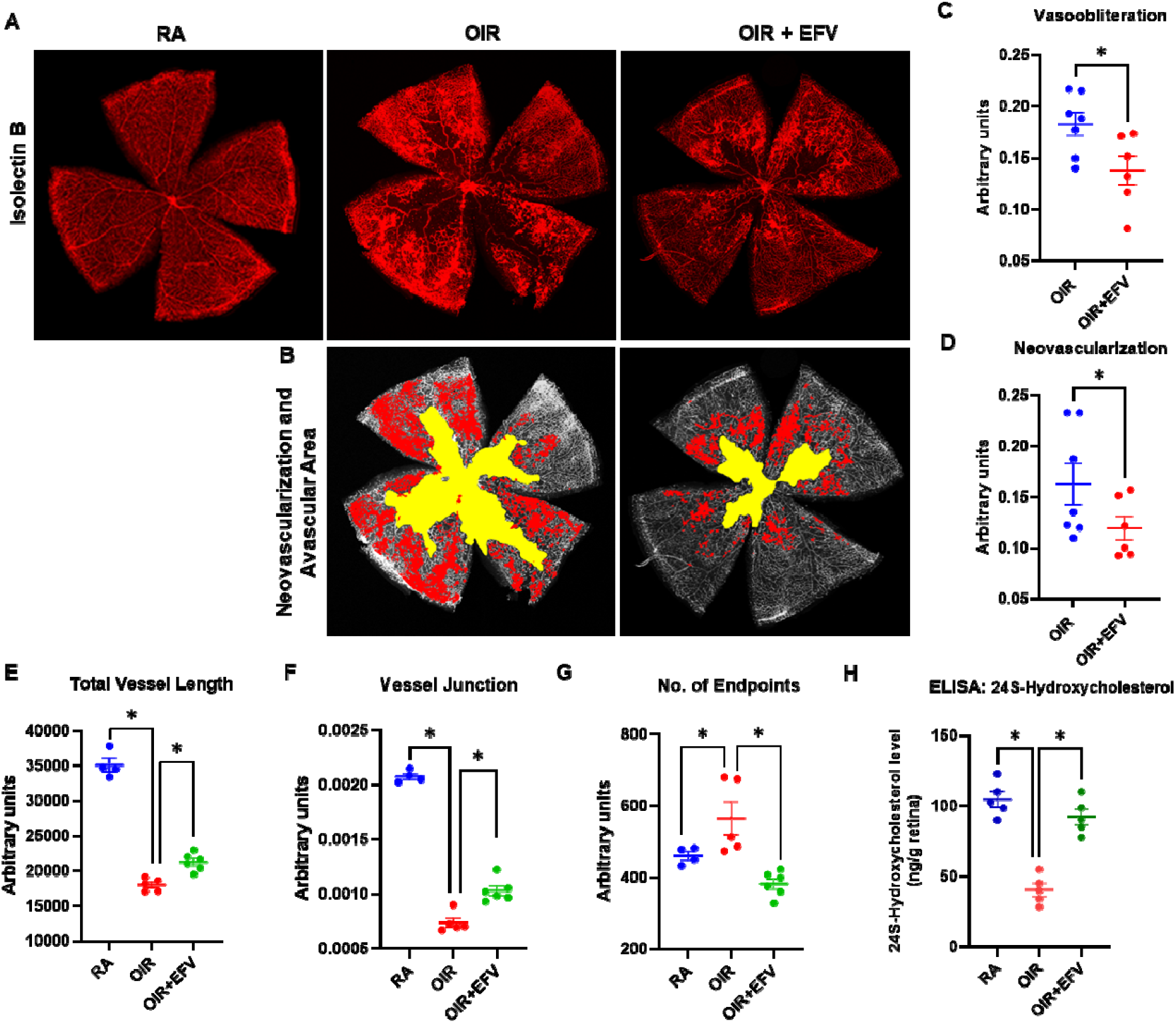
(A) Representative isolectin B4-stained retinal flatmounts prepared from room air (RA), oxygen-induced retinopathy (OIR), and efavirenz (EFV)-treated OIR (20 mg/kg; P7-P17) sacrificed at P17. (B) Representative retinal flatmounts in which the area of neovascularization (red) and avascular area (yellow) have been highlighted. Retinal images were further processed using OIRSeg, an unbiased automated web application, to quantitate changes in (C) vaso-obliteration (D) neovascularization. Further, vessel length (E), vessel junctions (F), and number of endpoints (G) were calculated using AngioTool analysis. Values are presented as mean ± SEM (n=4-7 retinas per group, each obtained from a different mouse). *p<0.05 vs. RA or OIR. (H) An ELISA was performed on retinal samples to compare levels of 24S-Hydroxycholesterol after activation of CYP46A1 by EFV treatment. Values are presented as mean ± SEM (n=5 retinas per group). *p<0.05 vs. RA or OIR.

Immunofluorescence analyses revealed diffuse CYP46A1 expression throughout the developing inner retina (data not shown). To better understand the cell-type specific expression of CYP46A1, Western blot analyses were performed using protein isolated from human retinal astrocytes (HRA), human brain microglia (HMC3), human retinal endothelial cells (HREC), human retinal pigment epithelial cells (ARPE), and differentiated retinal neuronal cells (RNC), that are all key in OIR pathology [31] (Fig. 4A). CYP46A1 expression was detected in each cell type. To better understand the mechanisms responsible for the improvements in OIR pathology realized in the presence of EFV/increased CYP46A1 expression atop the impact on retinal endothelial cells/blood vessels, we further investigated the response of macro-(Müller cells) and micro-glial cells of the retina as well as retinal ganglion cell viability. Müller cells exhibit reactive gliosis upon activation by inflammatory stimuli, marked by increased expression of GFAP [32]. This gliotic response of Müller cells is associated with the promotion of RNV and the breakdown of the blood-retinal barrier [33, 34]. Increased GFAP staining in the retina of OIR mice indicates gliosis, reflecting the activation of astrocytes and Müller cells in response to disease processes (Fig. 4B). EFV treatment significantly reduced gliosis in OIR mouse retinas as indicated by reduced GFAP immunoreactivity in astrocytes and Müller cells in EFV-treated compared to non-treated OIR mice (Fig. 4B). Müller cells communicate directly with retinal neurons, including retinal ganglion cells, neurons whose axons converge to form the optic nerve and are therefore critical to retinal function. Immunofluorescence analyses using the specific neural ganglion cell marker RBPMS [35], indicate that EFV treatment protected against retinal ganglion cell death in OIR mouse retinas (Fig. 4C).

**Figure 4.**
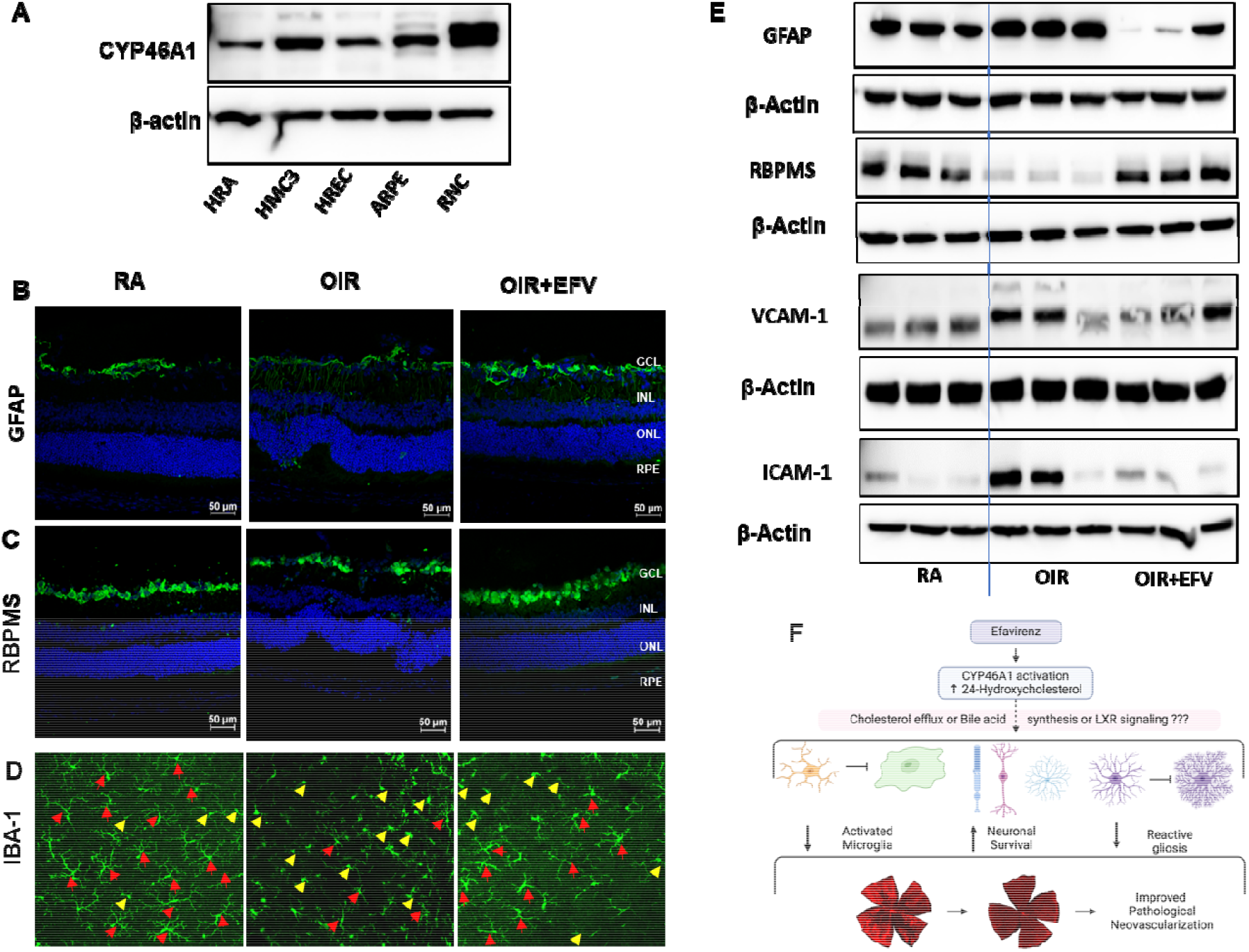
(A) Western blot analyses were performed using protein isolated from human retinal astrocytes (HRA), human brain microglia (HMC3), human retinal endothelial cells (HREC), human retinal pigment epithelial cells (ARPE), and differentiated retinal neuronal cells (RNC). Immunofluorescence staining of retinal cross sections from each experimental group to evaluate (B) GFAP and (C) RBPMS expression at P17. (D) IBA-1 immunostaining in representative retinal flat mount images from P17 mice of each group to identify quiescent (red arrowheads) or activated (yellow arrowheads) retinal microglia. (E) Western blot analyses of GFAP, RBPMS, VCAM-1 and ICAM-1 expression. β-actin was used as a loading control. Representative data presented in this figure are derived from 3-4 retinas per group, each obtained from a different mouse. (F) A proposed mechanism for CYP46A1-activation medicated effects of pathological neovascularization in mice.

Microglia directly guide vascular growth by interacting with the filopodia of endothelial tip cells during retinal vasculature formation [36]. Notably, in ischemic retinas, microglia tend to aggregate in ischemic and neovascularization regions, which are implicated as key contributors to the pathological processes. Infiltrating immune cells and local inflammatory signals can activate microglia. Activated microglia, in turn, produce cytokines and chemokines that further upregulate VCAM-1 and ICAM-1 expression, creating a vicious cycle of inflammation [37]. The inflammatory milieu created by activated microglia and infiltrating immune cells can disrupt the normal angiogenic process. This disruption can lead to the aberrant growth of retinal blood vessels characteristic of the later stages of ROP. Retinal flat mounts from RA, OIR, and OIR+EFV treated animals were subjected to immunofluorescence analyses of IBA-1 to facilitate the identification of ramified (quiescent) and amoeboid (activated) microglia (Fig. 4D). Ramified microglia were predominant in RA retinal flat mounts; however, flat mounts prepared from OIR mice were riddled with amoeboid microglia. Congruent with observations above, EFV treatment significantly reduced microglia activation which is associated with reduced inflammation in OIR mouse retinas (Figure. 4D). Western blot analyses confirm immunofluorescence staining results wherein EFV treatment significantly limited inflammation, ameliorated reactive gliosis and prevented neuronal cell loss in EFV-treated OIR mice (Fig. 4E).

## 4. Conclusions

The cytochrome P450 enzyme CYP46A1 is integral to cholesterol metabolism. Deficiency in CYP46A1 expression and activity in the adult mammalian retina is associated with impaired cholesterol metabolism and retinal morphology and function. The current study is the first, to our knowledge, to demonstrate the developmental regulation of CYP46A1 expression and the impact of its maintenance in the prevention and treatment of pathology in OIR mice, a model that mimics key mechanistic and pathological aspects of ROP in humans. Although, the precise mechanism(s) responsible for the observed CYP46A1-mediated effects, whether through enhanced cholesterol elimination, retinal bile acid signaling, or the LXR pathway, warrant further investigation (Fig. 4F), this study provides essential initial evidence that improving retinal cholesterol metabolism via CYP46A1 activation could offer therapeutic benefits for OIR mice and underscores the potential of targeting cholesterol metabolism in the treatment of ROP.

## Credit authorship contribution statement

**Briah Bailey and Josephine Rudd Zhong Manis:** Data curation, writing—original draft, methodology. **Allston Oxenrider, Gayatri Seth, and Shubhra Rajpurohit**: methodology and data curation. **Pamela M. Martin:** resources, supervision, and writing—review and editing. **Ravirajsinh N. Jadeja:** resources, validation, methodology, and writing—review and editing. **Menaka C. Thounaojam:** resources, supervision, project administration, funding acquisition, and writing—review and editing

## Funding

The realization of the proposed studies was made possible thanks to NIH grants, NIH EY34568 Grant (to M.C.T.); start□up funds from Augusta University (to M.C.T.) provided funding for this project. Additionally, we would like to express our appreciation to VDI AU Core Facilities for their support through the NIH EY031631 Grant. Student trainees were supported by GM144927, HL007737 and AI007281.

## Institutional Review Board Statement

All animal procedures were performed per protocols approved by Institutional Animal Care and Use Committee.

## Data Availability Statement

The original contributions presented in the study are included in the article/supplementary material; further inquiries can be directed to the corresponding author.

## Acknowledgments

We thank Dr. Syed Adeel Zaidi (AU) for providing proteins from differentiated retinal neuronal cells.

## Conflicts of Interest

The authors declare no conflicts of interest.

## Notes

### Competing Interest Statement

The authors have declared no competing interest.

### Summary of Updates

This version has been revised to update statistical analysis, representation of graphs, and some minor language edits.

## References

1. Casteels, I., et al., Educational paper: Retinopathy of prematurity. Eur J Pediatr, 2012. 171(6): p. 887–93.

2. Cavallaro, G., et al., The pathophysiology of retinopathy of prematurity: an update of previous and recent knowledge. Acta Ophthalmol, 2014. 92(1): p. 2–20.

3. Shah, P.K., et al., Retinopathy of prematurity: Past, present and future. World J Clin Pediatr, 2016. 5(1): p. 35–46.

4. Falavarjani, K.G. and Q.D. Nguyen, Adverse events and complications associated with intravitreal injection of anti-VEGF agents: a review of literature. Eye (Lond), 2013. 27(7): p. 787–94.

5. Hartnett, M.E. and J.S. Penn, Mechanisms and management of retinopathy of prematurity. N Engl J Med, 2012. 367(26): p. 2515–26.

6. Mutlu, F.M. and S.U. Sarici, Treatment of retinopathy of prematurity: a review of conventional and promising new therapeutic options. Int J Ophthalmol, 2013. 6(2): p. 228–36.

7. Hartnett, M.E., et al., Glaucoma as a cause of poor vision in severe retinopathy of prematurity. Graefes Arch Clin Exp Ophthalmol, 1993. 231(8): p. 433–8.

8. Choi, M.Y., I.K. Park, and Y.S. Yu, Long term refractive outcome in eyes of preterm infants with and without retinopathy of prematurity: comparison of keratometric value, axial length, anterior chamber depth, and lens thickness. Br J Ophthalmol, 2000. 84(2): p. 138–43.

9. Knight-Nanan, D.M., et al., Advanced cicatricial retinopathy of prematurity--outcome and complications. Br J Ophthalmol, 1996. 80(4): p. 343–5.

10. Kaiser, R.S., et al., Adult retinopathy of prematurity: outcomes of rhegmatogenous retinal detachments and retinal tears. Ophthalmology, 2001. 108(9): p. 1647–53.

11. Thounaojam, M.C., et al., Ursodeoxycholic Acid Halts Pathological Neovascularization in a Mouse Model of Oxygen-Induced Retinopathy. J Clin Med, 2020. 9(6).

12. Warden, C. and M.A. Brantley, Jr., Glycine-Conjugated Bile Acids Protect RPE Tight Junctions against Oxidative Stress and Inhibit Choroidal Endothelial Cell Angiogenesis In Vitro. Biomolecules, 2021. 11(5).

13. Win, A., et al., Pharmacological and Metabolic Significance of Bile Acids in Retinal Diseases. Biomolecules, 2021. 11(2).

14. Gong, Y., et al., Cytochrome P450 Oxidase 2C Inhibition Adds to omega-3 Long-Chain Polyunsaturated Fatty Acids Protection Against Retinal and Choroidal Neovascularization. Arterioscler Thromb Vasc Biol, 2016. 36(9): p. 1919–27.

15. Gong, Y., et al., Fenofibrate Inhibits Cytochrome P450 Epoxygenase 2C Activity to Suppress Pathological Ocular Angiogenesis. EBioMedicine, 2016. 13: p. 201–211.

16. Liao, W.L., et al., Quantification of cholesterol-metabolizing P450s CYP27A1 and CYP46A1 in neural tissues reveals a lack of enzyme-product correlations in human retina but not human brain. J Proteome Res, 2011. 10(1): p. 241–8.

17. Omarova, S., et al., Abnormal vascularization in mouse retina with dysregulated retinal cholesterol homeostasis. J Clin Invest, 2012. 122(8): p. 3012–23.

18. Zheng, W., et al., Spatial distribution of the pathways of cholesterol homeostasis in human retina. PLoS One, 2012. 7(5): p. e37926.

19. Saadane, A., et al., Retinal and nonocular abnormalities in Cyp27a1(-/-)Cyp46a1(-/-) mice with dysfunctional metabolism of cholesterol. Am J Pathol, 2014. 184(9): p. 2403–19.

20. Mast, N., et al., Cholestenoic Acid is an important elimination product of cholesterol in the retina: comparison of retinal cholesterol metabolism with that in the brain. Invest Ophthalmol Vis Sci, 2011. 52(1): p. 594–603.

21. Janowski, B.A., et al., An oxysterol signalling pathway mediated by the nuclear receptor LXR alpha. Nature, 1996. 383(6602): p. 728–31.

22. Thounaojam, M., et al., Differential bile acids biosynthetic pathways in the developing retina and in the oxygen-induced model of retinopathy of prematurity. Investigative Ophthalmology & Visual Science, 2021. 62(8): p. 3029–3029.

23. Gutsaeva, D.R., et al., STAT3-mediated activation of miR-21 is involved in down-regulation of TIMP3 and neovascularization in the ischemic retina. Oncotarget, 2017. 8(61): p. 103568–103580.

24. Smith, L.E., et al., Oxygen-induced retinopathy in the mouse. Invest Ophthalmol Vis Sci, 1994. 35(1): p. 101–11.

25. Pikuleva, I.A. and C.A. Curcio, Cholesterol in the retina: the best is yet to come. Prog Retin Eye Res, 2014. 41: p. 64–89.

26. Fliesler, S.J. and L. Bretillon, The ins and outs of cholesterol in the vertebrate retina. J Lipid Res, 2010. 51(12): p. 3399–413.

27. Saadane, A., et al., Retinal Vascular Abnormalities and Microglia Activation in Mice with Deficiency in Cytochrome P450 46A1-Mediated Cholesterol Removal. Am J Pathol, 2019. 189(2): p. 405–425.

28. Petrov, A.M., et al., CYP46A1 Activation by Efavirenz Leads to Behavioral Improvement without Significant Changes in Amyloid Plaque Load in the Brain of 5XFAD Mice. Neurotherapeutics, 2019. 16(3): p. 710–724.

29. Anderson, K.W., et al., Mapping of the Allosteric Site in Cholesterol Hydroxylase CYP46A1 for Efavirenz, a Drug That Stimulates Enzyme Activity. J Biol Chem, 2016. 291(22): p. 11876–86.

30. Moller, M., J. Fourie, and B.H. Harvey, Efavirenz exposure, alone and in combination with known drugs of abuse, engenders addictive-like bio-behavioural changes in rats. Sci Rep, 2018. 8(1): p. 12837.

31. Dai, C., et al., Neurovascular abnormalities in retinopathy of prematurity and emerging therapies. J Mol Med (Berl), 2022. 100(6): p. 817–828.

32. Bringmann, A., et al., Muller cells in the healthy and diseased retina. Prog Retin Eye Res, 2006. 25(4): p. 397–424.

33. Lundkvist, A., et al., Under stress, the absence of intermediate filaments from Muller cells in the retina has structural and functional consequences. J Cell Sci, 2004. 117(Pt 16): p. 3481–8.

34. Bai, Y., et al., Muller cell-derived VEGF is a significant contributor to retinal neovascularization. J Pathol, 2009. 219(4): p. 446–54.

35. Rodriguez, A.R., L.P. de Sevilla Muller, and N.C. Brecha, The RNA binding protein RBPMS is a selective marker of ganglion cells in the mammalian retina. J Comp Neurol, 2014. 522(6): p. 1411–43.

36. Checchin, D., et al., Potential role of microglia in retinal blood vessel formation. Invest Ophthalmol Vis Sci, 2006. 47(8): p. 3595–602.

37. Zhou, Z., et al., Distinguished Functions of Microglia in the Two Stages of Oxygen-Induced Retinopathy: A Novel Target in the Treatment of Ischemic Retinopathy. Life (Basel), 2022. 12(10).

